# Leveraging the Mendelian Disorders of the Epigenetic Machinery to Systematically Map Functional Epigenetic Variation

**DOI:** 10.1101/2020.11.08.373456

**Authors:** Teresa R. Luperchio, Leandros Boukas, Li Zhang, Genay O. Pilarowski, Jenny Jiang, Allison Kalinousky, Kasper D. Hansen, Hans T. Bjornsson

**Affiliations:** Department of Genetic Medicine, Johns Hopkins University School of Medicine, Baltimore, MD, USA; Department of Biostatistics, Johns Hopkins Bloomberg School of Public Health, Baltimore, MD, USA; Faculty of Medicine, School of Health Sciences, University of Iceland, Reykjavik, Iceland; Landspitali University Hospital, Reykjavik, Iceland

**Keywords:** histone machinery, immune dysfunction, chromatin, histone modification, epigenetics, Mendelian disorders, Kabuki syndrome, Rubinstein-Taybi syndrome

## Abstract

The Mendelian Disorders of the Epigenetic Machinery (MDEMs) have emerged as a class of Mendelian disorders caused by loss-of-function variants in epigenetic regulators. Although each MDEM has a different causative gene, they exhibit several overlapping disease manifestations. Here, we hypothesize that this phenotypic convergence is a consequence of common abnormalities at the epigenomic level, which directly or indirectly lead to downstream convergence at the transcriptomic level. Therefore, we seek to identify abnormalities shared across multiple MDEMs, in order to pinpoint locations where epigenetic variation is causally related to disease phenotypes. To this end, we perform a comprehensive interrogation of chromatin (ATAC-Seq) and expression (RNA-Seq) states in B cells from mouse models of three MDEMs (Kabuki types 1&2 and Rubinstein-Taybi syndromes). We build on recent work in covariate-powered multiple testing to develop a new approach for the overlap analysis, which enables us to find extensive overlap primarily localized in gene promoters. We show that disruption of chromatin accessibility at promoters often leads to disruption of downstream gene expression, and identify 463 loci and 249 genes with shared disruption across all three MDEMs. As an example of how widespread dysregulation leads to specific phenotypes, we show that subtle expression alterations of multiple, IgA-relevant genes, collectively contribute to IgA deficiency in KS1 and RT1. In contrast, we predict that KS2 does not have IgA deficiency, and confirm this observation *in vivo*. We propose that the joint study of MDEMs offers a principled approach for systematically mapping functional epigenetic variation in mammals.

## Introduction

A long-standing and fundamental problem in epigenetics is the identification of specific epigenetic changes that *causally* mediate phenotypes through the alteration of transcriptional states. While statistical associations between many diseases/traits and epigenetic changes have been detected, it is typically extremely challenging to rule out the influence of confounders such as the environment, and to determine whether these associations are primary causes versus secondary consequences^1,2^. As a result, to date there are surprisingly few examples of causal relationships between epigenetic alterations and specific phenotypes; notable exceptions include disorders of genomic imprinting^3^, disorders caused by repeat-expansion-induced aberrant promoter hypermethylation^4,5^, and metastable epialleles^6,7^.

The recent advent and widespread clinical use of exome sequencing has led to the emergence of a novel class of Mendelian disorders, termed the Mendelian Disorders of the Epigenetic Machinery (MDEMs)^8^. MDEMs are caused by coding variants disrupting genes encoding for epigenetic regulators, which are generally very intolerant to loss-of-function variation^9^. This implies the following causal chain underlying MDEM pathogenesis: a coding variant disrupts an epigenetic regulator, leading to downstream epigenomic abnormalities, which in turn give rise to the phenotype, likely by perturbing the transcriptome (**Figure 1a**). As a result, MDEMs may provide a unique lens into the causal relationship between epigenetic/transcriptomic variation and disease. Indeed, studies of Kabuki syndrome type 1 - one of the most extensively studied MDEMs to date - have begun unraveling the underpinnings of the neural^10^, growth^11^, cardiac^12^, and immune defects^13–15^ seen in this disorder.

**Figure 1.**
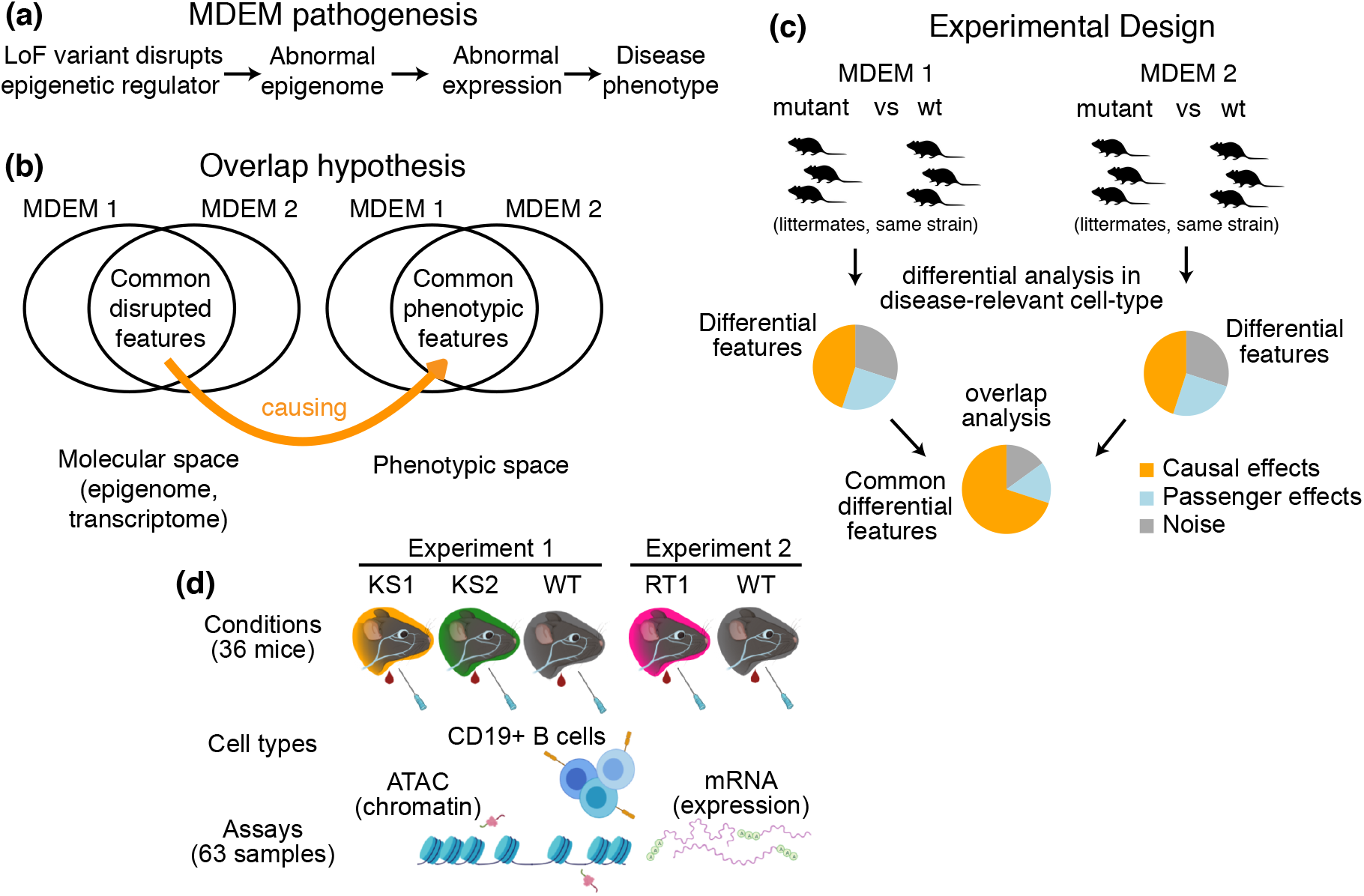
The theoretical framework for the present study. **a**) The causal chain of MDEM pathogenesis: the genetic disruption of an epigenetic regulator leads to epigenetic and transcriptomic consequences, which ultimately determine the phenotype. **b**) We hypothesize that the shared phenotypic features between MDEMs occur because of shared epigenetic and transcriptomic abnormalities downstream of the genetic disruption of distinct genes. The Venn diagram depicts two MDEMs for convenience, but our approach can be applied to an arbitrary number of MDEMs with shared phenotypes. **c**) Our approach is designed to derive a list of abnormalities with high probability of causal relevance, by jointly comparing multiple MDEMs. Shown for two MDEMs for convenience. **d**) Experimental design and workflow for sample generation in our present study. Created with BioRender.com

With studies of individual MDEMs, however, it remains challenging to distinguish the causal, disease-driving molecular alterations, from noise or passenger effects. As a consequence, it has been difficult to pinpoint specific target genes/loci that contribute to pathogenesis. It is also unclear if such targets span a small portion of the genome, or whether the disease phenotypes arise through the combined effects of multiple perturbations widely distributed across the genome.

## Results

### Joint analysis of multiple MDEMs to identify causally relevant epigenetic and transcriptomic variation

We set out to design an approach that discovers functionally relevant epigenetic variation, and overcomes limitations such as confounding effects from the environment and reverse causality from the disease process. Our approach leverages a cardinal and thus far unexploited feature of MDEMs, namely their overlapping phenotypic features, despite the causative genetic variants disrupting distinct genes. Such common MDEM features include intellectual disability, growth defects, and immune dysfunction^8^. We hypothesized that the common phenotypes arise because the different primary genetic defects lead to common downstream epigenomic alterations, which in turn create common transcriptomic alterations (**Figure 1b**). This hypothesis of a convergent pathogenesis motivates a joint analysis of more than one MDEM, and suggests a simple filter to identify the causal variation at the epigenetic/transcriptomic level: true, disease-driving signals should be detectable in multiple disorders (**Figure 1c**).

With this approach, two central practical considerations arise. First, what biological samples are most appropriate to allow for non-ambiguous interpretation of the results? We propose the use of MDEM mouse models, which have been shown to closely recapitulate many aspects of the human phenotypes^13,16,17^. Importantly, using mice allows us to: a) eliminate multiple confounders such as the environment, genetic background, age, and sex, and b) maintain a consistent sampling of disease-relevant cell types between individuals.

Second, what statistical approach should be employed for detection of the true common alterations? The simplest way would be to perform differential accessibility and expression analyses separately for each disorder, and obtain a list of the overlapping differential hits. However, this suffers from the major shortcoming that in order to be labeled as differential, a given locus must exceed an arbitrary significance threshold (or rank). When multiple MDEMs are studied, this requirement can lead to severe loss of power and erroneous underestimation of the size of the overlap among the differential hits. To avoid this, we recast the problem as testing whether evidence that a set of loci/genes are differential in a given MDEM is informative about the state (null or differential) of the same loci/genes in another MDEM. We show (**Methods**) that with this formulation, we can use conditional p-value distributions to: a) estimate the size of the set of overlapping abnormalities and test if it is greater than expected by chance, b) identify a set of genes that belong to this overlap, and c) decouple a) from b), so that only the identification of specific genes is affected by the multiple testing burden.

Here, as proof-of-principle, we implement our proposed approach using mouse models of 3 MDEMs: two that were clinically indistinguishable prior to the discovery of the underlying genes (Kabuki syndrome types 1 and 2; KS1 and 2, caused by haploinsufficiency in histone methyltransferases *KMT2D* and *KDM6A*, respectively), and one that shares phenotypes but is clinically distinct (Rubinstein-Taybi type 1; RT1, caused by haploinsufficiency in histone acetyltransferase *CREBBP*). All three syndromes have been previously found to exhibit immune system dysfunction. In KS1, this includes combined variable immunodeficiency with low IgA, as well as abnormal cell maturation^8,18,19^. In RT1, it includes hypogammaglobulinemia with a reduction of mature B cells^20^, while in KS2 the immune phenotype has been less extensively studied, with some evidence of mild hypogammaglobulinemia^18,21^. Given this potential overlap, we chose to profile positively selected B cells (CD19+; **Methods**) from the peripheral blood of mutant mice, and that of age- and sex-matched wild-type littermates (**Figure 1d**).

### Genome-wide chromatin accessibility profiling reveals extensive overlap between the epigenetic aberrations of the three MDEMs

We first used ATAC-seq to profile genome-wide chromatin accessibility, employing a modified fastATAC protocol (**Methods**)^22,23^. Starting with a differential accessibility analysis of 7 KS1 versus 12 wild-type mice (**Methods**), we discovered 3,121 ATAC peaks differentially accessible at the 10% FDR level. Of these, 824 (26.4%) overlapped promoters (defined as +/- 2kb from the TSS), and 2,297 (74%) were in distal regulatory elements (defined as ATAC peaks outside of promoters).

We then compared KS1 to KS2, focusing on promoters first. Using our new approach to detect overlap between lists of differential features from separate differential analyses (**Methods**), we found that 71.5% of promoter peaks differentially accessible in KS1 are also differential in KS2 (**Figure 2a**; p < 2.2e-16, 5 KS2 vs 12 wild-type mice); at the 10% FDR level, we identified 645 such peaks (**Supplemental Table 1**). For 630 of the 645 (97.7%), accessibility is altered in the same direction in the two syndromes (**Figure 2b**). Out of the 630 promoter peaks disrupted in both KS1 and KS2, we discovered that 62.2% are differential in RT1 as well (**Figure 2c**; p = 0.0034, 5 RT1 vs 7 wild-type mice), again with highly concordant effect sizes (**Figure 2d**).

**Figure 2.**
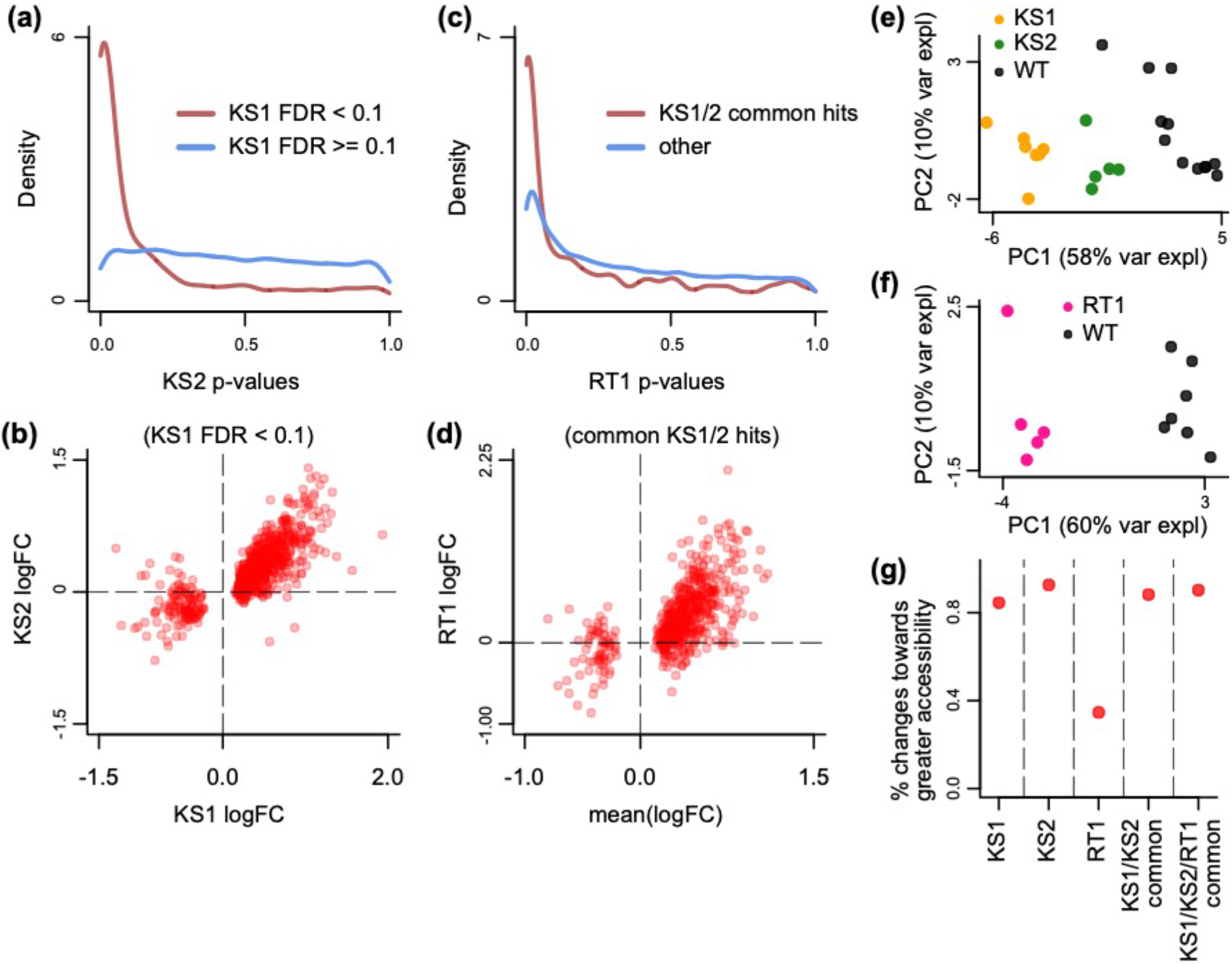
Evaluating the overlap between the differentially accessible promoter peaks in B cells in Kabuki type 1, Kabuki type 2, and Rubinstein-Taybi syndromes. **a**) The distribution of p-values from the KS2 vs wild-type differential accessibility analysis for promoter peaks, stratified according to whether the same promoter peaks are significantly differentially accessible in the KS1 vs wild-type analysis (FDR < 0.1; red curve), or not (FDR >= 0.1; blue curve). **b**) Scatterplot of log2(fold changes) from the KS1 vs wild-type (x-axis) promoter peak differential accessibility analysis against the corresponding log2(fold changes) from the KS2 vs wild-type analysis (y-axis). Each point corresponds to a peak. Shown are only peaks that are differentially accessible in KS1 (FDR < 0.1). **c**) The distribution of p-values from the RT1 vs wild-type differential accessibility analysis for promoter peaks, stratified according to whether the same promoter peaks are commonly differentially accessible in KS1 and KS2 (FDR < 0.1, see Methods; red curve), or not (blue curve). **d**) Scatterplot of log2(fold changes) from the RT1 vs WT (x-axis) differential accessibility analysis, against the mean log2(fold change) from the KS1 vs wild-type and KS2 vs wild-type analyses. Each point corresponds to a peak. Shown are only peaks that are commonly differentially accessible in KS1 and KS2 (FDR < 0.1). **e**) and **f**) Principal component analysis plots using only the 313 promoter peaks identified as commonly differentially accessible between the three MDEMs. Each point corresponds to a mouse. **g**) The proportion of differentially accessible promoter peaks that show increased accessibility in the mutants vs the wild-type mice.

In total, we identified 313 promoter peaks that show disruption in all 3 disorders at the 10% FDR level (**Supplemental Table 2**). This is ~4 times more shared peaks than we find if we perform separate differential analyses and compute the intersection of the resulting differential hits (78 peaks), highlighting that our new approach provides substantial gain in empirical power. A principal component analysis shows that the accessibility signal of these commonly disrupted promoter peaks separates each of the three mutant genotypes from their wild-type littermates (**Figure 2e, f**). This is most surprising for the case of KS1 and KS2, patients of which have such strong phenotypic overlap that the disorders were not considered distinct prior to discovery of the causative genes (**Figure 2e**).

Next, we applied the same approach to distal regulatory elements. We saw a similar picture, albeit with weaker shared signal (**Supplemental Figures 1a-d**). Specifically, 55.6% of elements differential in KS1 were estimated as differential in KS2 (p < 2.2e-16), with 958 confidently labeled such elements (10% FDR; **Supplemental Table 3**). As with promoters, we observed agreement in directionality. Of these 958 elements, 35.4% are shared with RT1 (p = 0.0053), yielding a total of 150 commonly disrupted distal regulatory elements across the 3 MDEMs (10% FDR; **Supplemental Table 4**). We note that, collectively, the common hits show a 5.3-fold enrichment at promoters compared to distal elements (Fisher’s test, p < 2.2e-16).

Finally, comparing the 3 MDEMs in a pairwise fashion, we observed that KS1 and KS2 share a greater proportion of their abnormalities than either KS1 or KS2 compared to RT1 (**Supplemental Figure 1e**), and verified that this is not driven by the fact that the KS1 and KS2 mice were compared against the same wild-type group (**Methods**).

### Shared disrupted promoters, but not distal regulatory elements, show bias towards increased accessibility in KS1 and KS2

We explored the direction in which the accessibility of the disrupted peaks changes in mutants compared to wild-type. We found that, at promoters, both the KS1 and KS2 mutants exhibit a substantial shift towards increased accessibility (84.6% and 92.7%, respectively, of significantly disrupted promoter peaks; **Figure 2g**). The same shift is observed in the promoter peaks commonly disrupted across the MDEMs (**Figure 2g**), even though in the RT1 mutants the majority (65.4%) of differentially accessible promoter peaks show the opposite pattern, with a shift towards decreased accessibility (**Figure 2g**). In contrast to promoters, disrupted distal regulatory elements in all cases are split: about half show increased accessibility and half show decreased accessibility (**Supplemental Figure 1f**).

### Transcriptome profiling reveals many expression alterations at genes downstream of promoters with disrupted accessibility

We next interrogated the transcriptome using RNA-seq (**Methods**) to: a) test whether the identified epigenetic aberrations in each disorder have direct transcriptional consequences and characterize the latter, and b) identify the shared expression aberrations across the three disorders, and assess the extent to which these result from shared accessibility aberrations at the associated promoters. To better understand the relationship between chromatin accessibility and gene expression, we generated the RNA-seq samples in parallel with the samples used for ATAC-seq, from a subset of the same individual mice; this allowed us to capture both chromatin and transcriptional status at a single time point.

First, for each disorder, we determined the top differential promoter peaks as ranked by p-value, and estimated the percent of genes downstream of these promoters that show differential expression; we repeated this by sliding the threshold for inclusion of the top ranked peaks from 1000 to 5000. When considering the top 1000 promoter peaks, the percentage of differentially expressed downstream genes is 36.2% in KS1, 41.3% in KS2, and in 49.5% in RT1 (**Methods**; 5 mice per genotype; **Supplemental Table 5** contains such genes detected at the 10% FDR level). In all three syndromes, this percentage gradually drops substantially as the cutoff for labeling a promoter as differentially accessible becomes less stringent (**Figure 3a**), indicating a clear relationship between abnormal promoter accessibility and downstream gene expression dysregulation. Emphasizing this relationship, we discovered strong concordance between the direction of abnormal changes at the disrupted promoter-gene pairs: increased or decreased promoter accessibility correlates with increased or decreased gene expression, respectively (**Figure 3b, c, d**; Pearson correlation between promoter accessibility logFC and gene expression logFC = 0.67 for KS1, 0.79 for KS2, and 0.82 for RT1).

**Figure 3.**
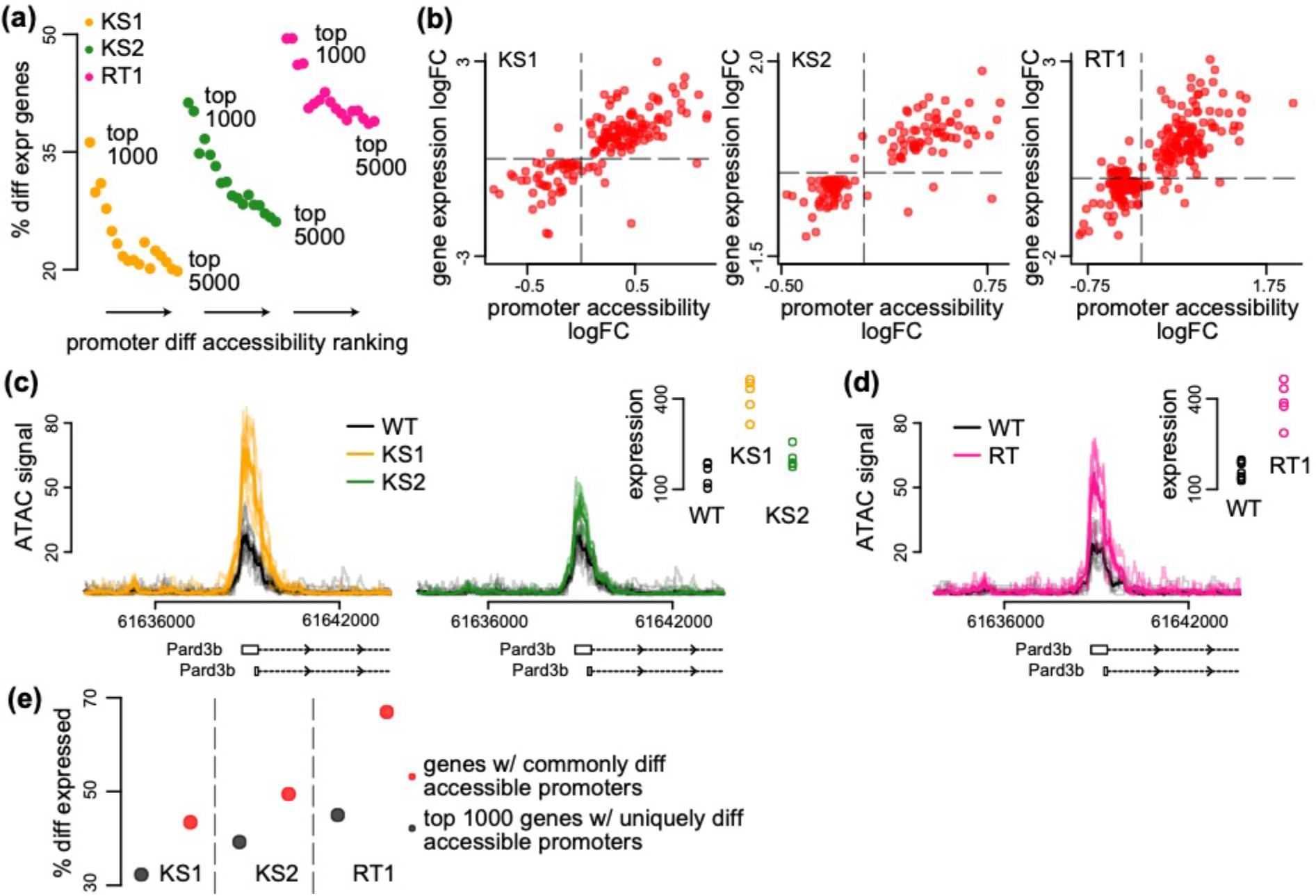
The relationship between differential accessibility of promoter peaks and differential expression of downstream genes in the three MDEMs. **a**) The proportion of promoters with differentially expressed downstream genes in KS1, KS2, and RT, estimated for the top ranked differentially accessible promoter peaks. The estimation was repeated for different thresholds for inclusion into the top ranked list. For each MDEM, each point corresponds to a different threshold. Thresholds were slid from 1000 to 5000, in steps of 250. **b**) Scatterplot of the accessibility log2(fold changes) of differentially accessible promoter peaks, against the expression log2(fold changes) of differentially expressed downstream genes, for each of the three MDEMs. Shown are only pairs where the promoter peak was within the top 1000 differentially accessible promoter peaks (ranked based on p-value), and the downstream gene was differentially expressed (10% FDR; methods). Each point corresponds to a gene-promoter pair. In cases where more than 1 peak in the same promoter was within the top 1000 differentially accessible peaks, the median(log2(fold change)) across all such peaks was calculated. **c**) and **d**) An example locus (*Pard3b*) with concordant changes in promoter peak accessibility and downstream gene expression in all three MDEMs. **e**) The proportion of promoters with differentially expressed downstream genes in KS1, KS2, and RT1, estimated separately for the top 1000 uniquely differentially accessible promoters in each MDEM, versus the same proportion estimated for the genes downstream of the 313 commonly differentially accessible promoter peaks.

Finally, we compared the proportion of differentially expressed genes downstream of the shared disrupted promoter peaks, to the same proportion for genes downstream of the top 1000 disrupted promoter peaks unique for each disorder. We invariably found the genes downstream of the shared disrupted peaks to have a higher chance of dysregulated expression (**Figure 3e**), supporting our hypothesis that the chromatin alterations at these peaks are more likely to lead to functional effects on transcription.

### A substantial proportion of the shared expression aberrations among the three MDEMs arise without concomitant disruption of promoter accessibility

To further dissect the relationship between the common expression and chromatin abnormalities in the three MDEMs, we sought to define a set of genes commonly differentially expressed, without utilizing prior information about the accessibility of their promoter peaks.

Utilizing our method, we discovered high overlap between KS1 and KS2, mirroring the findings at the chromatin level (**Figure 4a, b**). Specifically, we found 372 differentially expressed genes shared between them with concordant direction of effect (10% FDR; **Supplemental Table 6**). We then estimated 70% of these to be differential in RT1 (**Figure 4c, d**), resulting in 249 genes shared across the 3 disorders, with a preponderance of downregulated genes (**Figure 4e, f, g; Supplemental Table 7**; 171 downregulated vs 78 upregulated genes).

**Figure 4.**
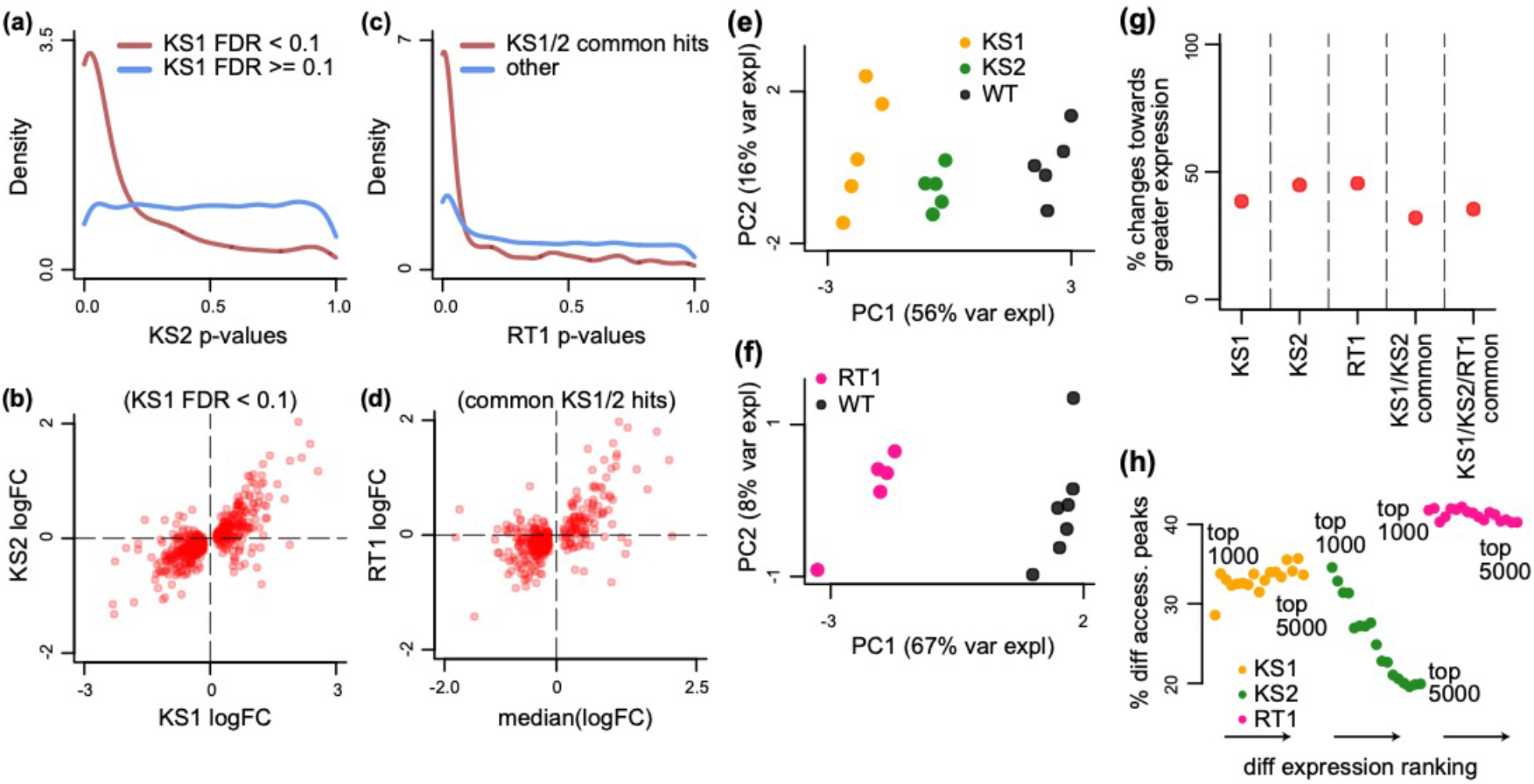
Evaluating the overlap between the differentially expressed genes in B cells in Kabuki type 1, Kabuki type 2, and Rubinstein-Taybi syndromes. **a**) The distribution of p-values from the KS2 vs wild-type differential expression analysis, stratified according to whether the same genes are significantly differentially expressed in KS1 (FDR < 0.1; red curve), or not (FDR >= 0.1; blue curve). **b**) Scatterplot of log2(fold changes) from the KS1 vs wild-type differential expression analysis (x-axis), against the corresponding log2(fold changes) from the KS2 vs wild-type analysis (y-axis). Each point corresponds to a gene. Shown are only genes that are differentially expressed in KS1 (FDR < 0.1). **c**) The distribution of p-values from the RT1 vs wild-type differential expression analysis, stratified according to whether the same genes are commonly differentially expressed in KS1 and KS2 (FDR < 0.1, see Methods; red curve), or not (blue curve). **d**) Scatterplot of log2(fold changes) from the RT1 vs WT (x-axis) differential expression analysis, against the mean log2(fold change) from the KS1 vs wild-type and KS2 vs wild-type analyses. Each point corresponds to a gene. Shown are only genes that are commonly differentially expressed in KS1 and KS2 (FDR < 0.1). **e**) and **f**) Principal component analysis plots using only the 249 genes identified as commonly differentially expressed between the three MDEMs. Each point corresponds to a mouse. **g**) The proportion of differentially expressed genes that show increased expression in the mutant vs the wild-type mice. **h**) The proportion of genes with differentially accessible promoter peaks in KS1, KS2, and RT, estimated for the top ranked differentially expressed genes. The estimation was repeated for different thresholds for inclusion into the top ranked list. For each MDEM, each point corresponds to a different threshold. Thresholds were varied from 1000 to 5000, in steps of 250.

While these 249 genes are significantly enriched in the set of genes with shared disruption of promoter accessibility (p=3.06e-5), the magnitude of this enrichment is modest (24 genes in the intersection; odds ratio = 2.79, **Table 1**). The number of such genes increases to 81 when we also include those harboring commonly disrupted regulatory elements nearby (+/- 1Mb from their promoter peaks). Taken together, these results indicate that there is convergent dysregulation at the expression level in these three MDEMs, which is downstream of the shared differential epigenetic alterations. Further underscoring this, we found that in KS1 and RT1 – but not in KS2 – the top differentially expressed genes are no more likely to have disrupted promoters than genes further down the differential list (**Figure 4h**).

**Table 1.**
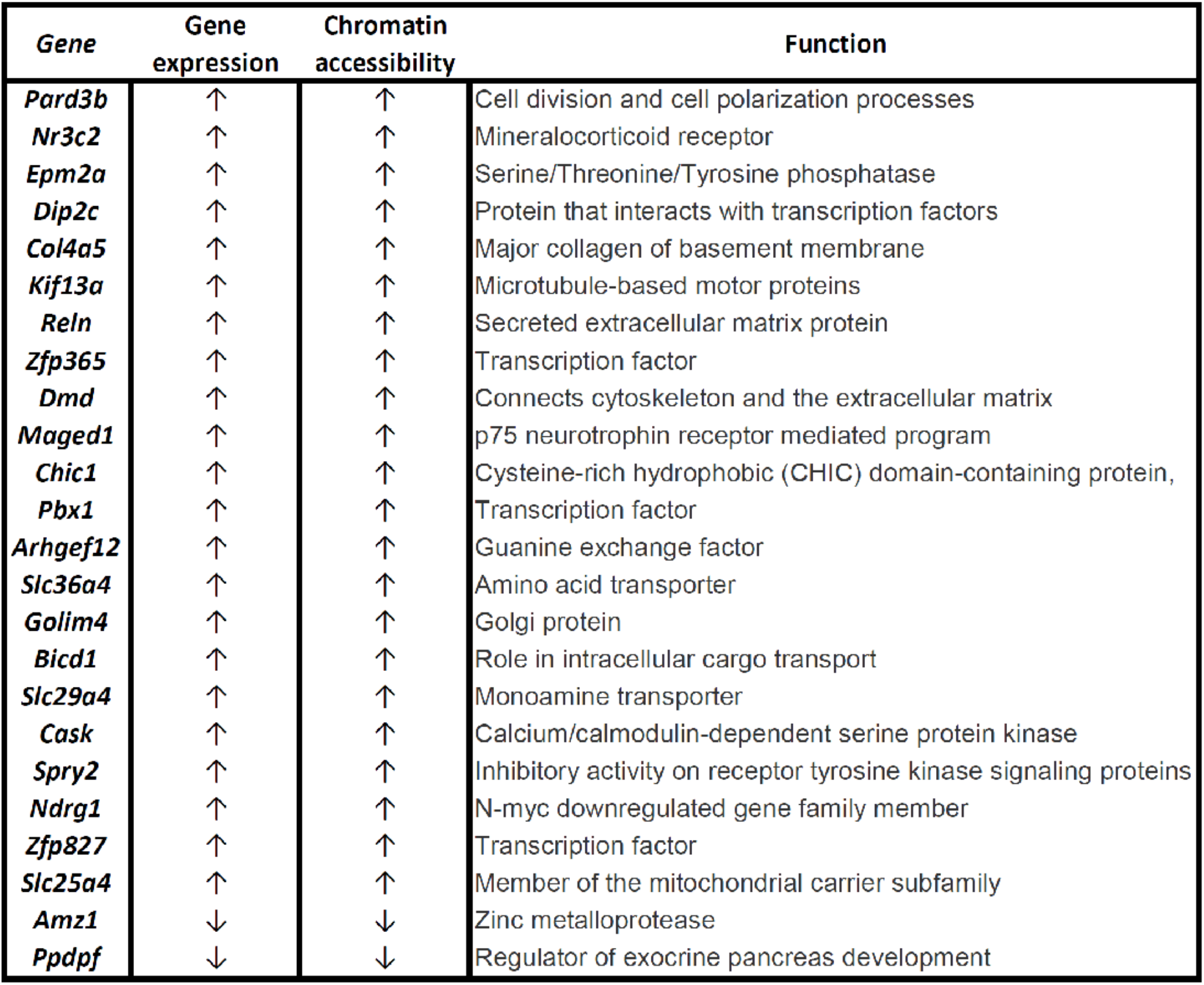

### Integration of transcription factor motifs with chromatin and expression abnormalities highlights some potentially disrupted regulatory connections

It is well-appreciated that chromatin accessibility is intimately linked to transcription factor binding. Accessibility patterns are often established subsequently to recruitment of epigenetic regulators by transcription factors at specific genomic locations^24^, while other transcription factors can only bind their cognate motifs if these reside within pre-accessible sites^25,26^. We therefore investigated the transcription factor motifs encoded within the differentially accessible peaks in the three disorders, using a set of 233 non-redundant motifs (**Methods**). We focused on differentially accessible peaks within promoters of differentially expressed genes, reasoning that these are more likely to reveal potentially disrupted regulatory connections with functional relevance. In all three syndromes, however, no single motif reached significant enrichment at the 10% FDR level when compared to other peaks.

Regulatory wiring disruption can occur not only because of altered motif accessibility, but also theoretically because of abnormal expression of the cognate transcription factors themselves. We thus also performed a search for motif enrichment in promoter peaks corresponding to differentially expressed genes, regardless of whether these peaks are differentially accessible or not, and then intersected the resulting motifs with our differentially expressed genes. The significant hits (**Supplemental Table 8**) included *Runx1*, as well as motifs recognized by transcription factors of the NF-kB pathway.

### The collective effect of individually subtle aberrations in multiple genes is likely responsible for perturbed IgA production and abnormal B-cell maturation

Our results establish the existence of widespread epigenetic and transcriptional aberrations that are largely shared across the three disorders, suggesting functional relevance. We therefore asked whether these aberrations can explain some specific aspects of the immune dysfunction. We first performed a pathway analysis of the shared disrupted genes (either at the expression or promoter accessibility level; **Methods**). This yielded several potentially affected pathways (**Supplemental Tables 9 and 10**). However, most of these were of general relevance and did not pinpoint very specific pathologies.

We then reasoned that we might gain more insight by focusing on two of the specific phenotypes seen in KS1: abnormal B-cell maturation, and IgA deficiency^13,20^. We set out to test if these are attributable to the collective dysregulation of multiple genes, or to the abnormal expression of a select few. To define relevant gene sets, we first obtained the set of all transcription factors encoded in the mouse genome that are expressed in CD19+ B cells (**Methods**); this choice was motivated by the fact that transcription factors are critical regulators of cellular differentiation and maturation. We then examined the ranks of these transcription factors in the KS1 p-value distribution, and observed a strong shift indicative of global dysregulation (**Figure 5a**; p < 2.2e-16). For IgA deficiency, we assembled a list of 75 genes known to lead to IgA deficiency when individually knocked out in mouse (**Methods**). Examination of the KS1 p-value ranks of these genes also highlighted a collective shift towards lower p-values (**Figure 5b**; p=0.02). Together, these results strongly suggest the collective (but often subtle) dysregulation of many genes.

**Figure 5.**
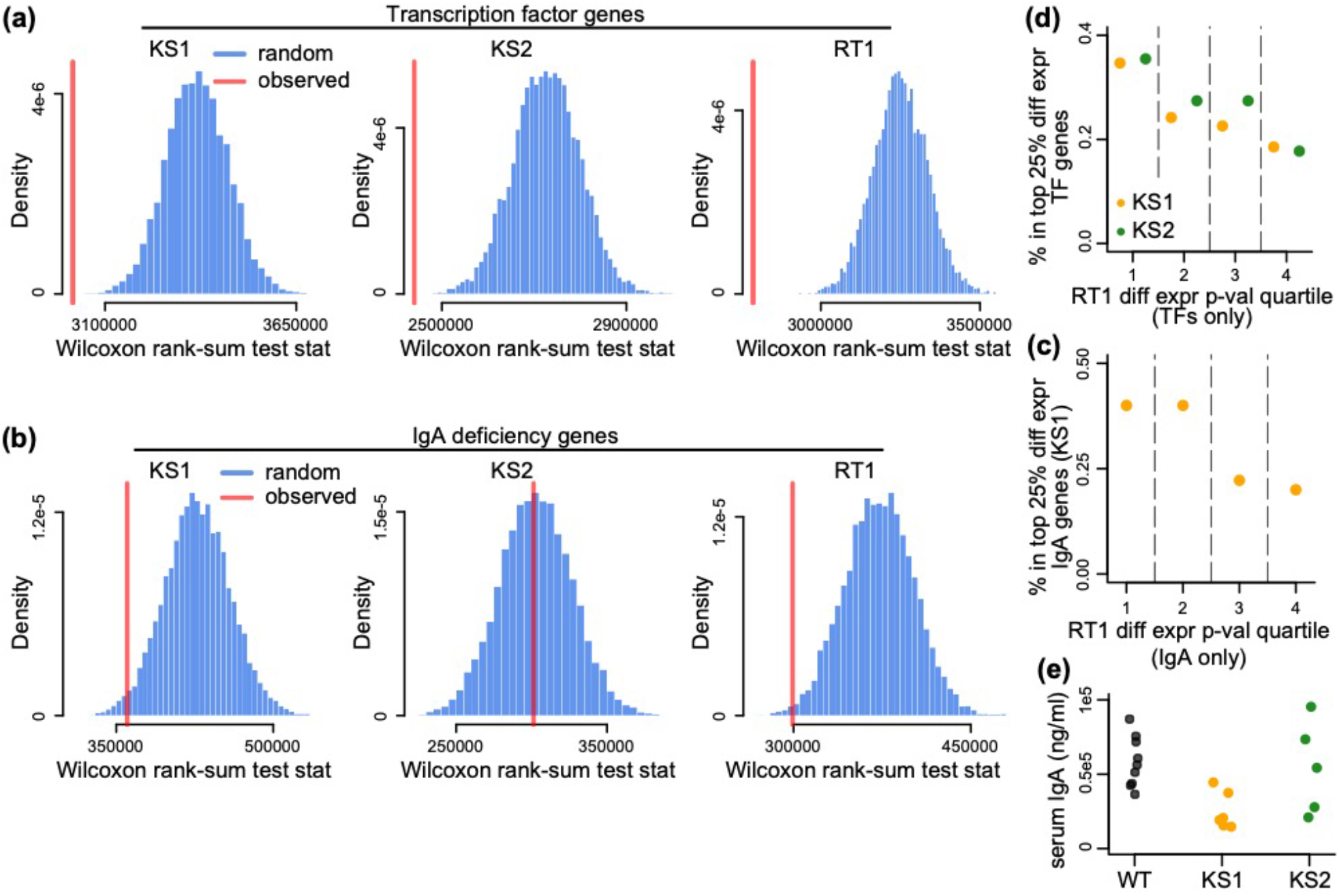
Evaluating genes known to encode for transcription factors, or individually contribute to IgA deficiency, for collective expression dysregulation. **a**) The Wilcoxon rank-sum test statistic (red vertical line) computed after assembling a list of genes encoding transcription factors expressed in B cells (Methods), and comparing the distribution of their differential expression p-values to the p-value distribution of the rest of the genes included in the differential expression analysis. The blue distribution corresponds to the same statistic computed after randomly sampling gene sets of the same size as transcription factors, and comparing their p-value distribution to the p-values for the rest of the genes. The resampling was performed 10,000 times. **b**) Same as **a**), but for genes known to individually contribute to IgA deficiency (Methods). **c**) The percentage of TF genes that belong to the top 25% differentially expressed TFs in KS1 (orange dots), and KS2 (green dots), stratified according to their p-value quartile in RT. **d**) Same as c), but for IgA deficiency genes compared in KS1 and RT. **e**) Serum IgA levels in KS1, KS2 and wild-type mice.

Turning our attention to KS2 and RT1, we observed similar results for transcription factors, with substantial contribution from a set of transcription factors dysregulated in all three MDEMs (**Figure 5a, c**). However, when assessing the IgA deficiency genes, we only observed the signal in RT1, and not in KS2 (**Figure 5b, d**). This was surprising, given the high phenotypic similarity between KS1 and KS2, and prompted us to measure serum IgA in the KS1/2 and wild-type mice (**Methods**). In agreement with the collective behavior of IgA-related genes, we found no difference in IgA levels between the KS2 and wild-type, while we recapitulated our previous result of IgA deficiency in KS1 mice^13^ (**Figure 5e**; p = 0.8 for KS2 vs WT, p = 0.0008 for KS1 vs WT).

Finally, we found no evidence that these collective defects in the expression of transcription factors and IgA-deficiency associated genes are driven by similar shifts towards abnormal promoter accessibility (p > 0.1 in all cases).

## Discussion

Our study shows that three MDEMs caused by loss-of-function variants in three distinct epigenetic regulators, share common abnormalities at the chromatin and gene expression level. These abnormalities show evidence of functionality, as illustrated by the fact that: a) many chromatin changes at promoters are linked to downstream gene expression changes, and b) systematic expression changes affect genes known to contribute to specific, well-characterized phenotypic features (IgA deficiency, abnormal B cell maturation) seen in these MDEMs.

In terms of understanding the pathogenesis of MDEMs, our results clearly point towards a generalized, systems-level dysregulation, with a multitude of cellular processes/pathways affected. From our present study it is unclear how exactly these combine to ultimately give rise to the phenotypic manifestations; elucidating this will be an important challenge going forward. It is also worth noting that the emergent picture bears similarities to the molecular basis of complex diseases, with many widely distributed, small-effect perturbations, ultimately generating the phenotype^27^. This is perhaps not unexpected, given that epigenetic regulators are typically *trans*-acting proteins that act at many locations. It also suggests that, even though MDEMs are single-gene Mendelian disorders, they might best be conceptualized as effectively complex disorders. This may also explain the broadness of the phenotype in MDEMs^28^, and the decreased penetrance of many phenotypes in patients, that are fully penetrant in mouse models.

It is notable that we find greater molecular overlap between KS1 and KS2 than between either of them and RT1, in agreement with the greater similarity between the two KS types at the phenotypic level. It should be mentioned, however, that specific sub-phenotypes provide exceptions to this rule, as evidenced by the multi-genic abnormalities in IgA deficiency genes, which are shared between KS1 and RT1 but are absent in KS2. Together, these results suggest that deep phenotyping of MDEMs combined with molecular characterization can yield new insights into the pattern of their shared features.

One unexpected finding was that, at promoters, almost all of the shared disrupted peaks exhibit a shift towards a more open chromatin state, even though the causative mutations of all three disorders would theoretically be expected to push towards a more closed chromatin state, based on the specific histone marks they are thought to affect^29^. One possible explanation is that these shared hits represent indirect effects, arising downstream of the initial effects of the mutations. Alternatively, the hypothesis that loss of the epigenetic regulators disrupted in our three disorders would lead to closed chromatin may not hold. Finally, there is the possibility that the causative mutations lead to a non-specific cellular compensatory response, which causes increased chromatin openness at several genomic locations such as the adaptive stress response^30^. The latter is supported by the fact that many of the shared genes uncovered in this study are not known to be direct KMT2D, KDM6A or CREBBP target genes. Regardless of the exact reason, this observation warrants future exploration.

We note that our study differs from recent studies of DNA methylation in the peripheral blood of MDEM patients. In these studies, the goal is to derive “episignatures” with the capacity for robust phenotypic prediction. While the predictive power of these episignatures is evidenced by their ability to provide a diagnosis^31^, their functional relevance is uncertain, for two main reasons. First, peripheral blood is a mixture of several cell types, and changes in cell-type composition - which often occur in disease - can severely confound the differential DNAm analysis^32,33^. Second, the great majority of these MDEMs are caused by variants in histone modifiers or chromatin remodelers. As a result, even if cell-type heterogeneity were to be ignored, these DNAm aberrations are by definition secondary events. In contrast, our strategy is specifically designed to yield a catalog of abnormalities with primary functional role in common MDEM pathogenesis.

In summary, we propose the study of the Mendelian Disorders of the Epigenetic Machinery as a principled approach for systematically mapping causally relevant epigenetic variation in mammals. The shared hits among the three disorders studied here almost exclusively demonstrate an increase in open chromatin at promoters, which is counterintuitive to the function of the individual causative genes and may either suggest a previously unexpected role for them, or an undescribed systemic compensatory response. Finally, we suggest that MDEMs are effectively complex disorders, arising from widely distributed epigenetic perturbations across the genome.

## Methods

### Mice

We performed all mouse experiments in accordance with the National Institutes of Health Guide for the Care and Use of Laboratory Animals and all were approved by the Animal Care and Use Committee of the Johns Hopkins University. We genotyped mice using standard genotyping and PCR methods. For all comparisons, we used wild type littermates.

*KS1. Kmt2d^+/βGeo^* mice are fully backcrossed to C57BL/6J and this backcrossing is verified by SNP genotyping^17^. These mice are also known as *Mll2^Gt(RRt024)Byg^*, and originally obtained from BayGenomics and fully backcrossed in the Bjornsson laboratory.

#### Primers

βGeo F-CAAATGGCGATTACCGTTGA, R-TGCCCAGTCATAGCCGAATA; Tcrd (control) F-CAAATGTTGCTTGTCTGGTG, R-GTCAGTCGAGTGCACAGTTT *KS2. Kdm6a^+/-^* mice were acquired from European Mouse Mutant Archive (EMMA) but this model has also been called: *Kdm6a^tm1a(EUCOMM)Wtsi^*. Mice were crossed with flippase expressing mice (B6.Cg-Tg(ACTFLPe)9205Dym/J) (Jackson Laboratories) to remove the third exon of *Kdm6a*, and then progeny were crossed with Cre expressing mice driven by CMV (B6.C-Tg(CMV-cre)1Cgn/J) (Jackson Laboratories), to generate the *Kdm6a^tm1d(EUCOMM)Wtsi^* allele. Mice were backcrossed on C57BL/6J to maintain the *Kdm6a^tm1d(EUCOMM)Wtsi^* allele.

#### Primers

Kdm6aTm1c F-AAGGCGCATAACGATACCAC, Floxed LR-ACTGATGGCGAGCTCAGACC; Tcrd (control) F-CAAATGTTGCTTGTCTGGTG, R-GTCAGTCGAGTGCACAGTTT *RT1. Crebbp^+/-^* mice also known as *Crebbp^tm1Dli^*, were acquired from Jackson laboratories but established by Kung et al^16^. These mice were maintained on a C57BL/6J background in the Bjornsson laboratory.

#### Primers

R-T F: TAAGCAGCAGCATCCTTTGG, R-T_WT R: CCTGACAATGTGTCATGTGAT, R_T_MUT R: ATGCTCCAGACTGCCTTGGGA;

### Sex disaggregation

We performed all experiments in female mice to enable comparison between all three disease models as *KDM6A* (KS2 model) is present on the X chromosome and is lethal in male mice when knocked out. Therefore, we are unable to present sex-disaggregated data.

### Blood cell isolation

We obtained peripheral blood from 2.5-3.5 month old female mice by facial vein bleed. 150-250 μl blood was collected in K2EDTA blood collection tubes (BD Microtainer 365974) and red blood cells were lysed for 7-15 minutes at room temperature in 2mL red blood cell lysis solution (15.5mM NH4Cl, 1mM KHCO3, 0.01mM EDTA). We diluted lysed blood with excess balanced salt solution (Gey’s or 1xPBS), manually removed large clots using pipet tip, and spun at 500g for 10 minutes 4’. Second lysis at room temperature was performed for samples with large amounts of remaining red blood cells then spun. We isolated CD19+ B cells by positive selection using CD19+ microbeads for mouse (Miltenyi 130-052-201) following manufacturer protocols, then counted and aliquoted samples on ice to further process for ATAC-seq and RNA-seq.

### ATAC-seq

We performed ATAC-seq using a modified FastATAC protocol^22,23^. Specifically, we resuspended 5k cells per reaction in 1xPBS and quickly spun to remove residual EDTA from isolation steps, and then resuspended in tagmentation reaction mix for 30 min (2.5uL TD1, 1X TD Buffer, Illumina Nextera DNA, FC-121-1030; .25uL 1% digitonin, Promega G9441; 1xPBS;) gently shaking (300rpm on Eppendorf thermomixer) at 37’. We purified reactions using Zymo DNA Clean and Concentrator-5 kit (Zymo D4013) following manufacturer protocols and eluted with 10.5μL water to recover 10μL. Each reaction was then amplified and indexed as described^22^, total sample amplification cycles range from 6-10 cycles. After indexing and amplification, we purified samples using Select-A-Size purification columns (Zymo D4080) with a cutoff of 150bp to remove adapter dimers to allow for efficient sequencing on patterned flow cells, checked library size on BioAnalyzer using DNA High Sensitivity reagents (Agilent 5067-4626) and determined concentration using Qubit dsDNA HS Assay Kit (ThermoFisher Q32851). We pooled and sequenced on Illumina HiSeq4000 using PE flow cells with 100-8-8-100 read length using standard manufacturer protocols. Samples were clustered to aim for 60M reads per sample Samples were demultiplexed using Illumina pipeline bcl2fastq2 v2.20 with all defaults except -- use-bases-mask Y100n, I8, I8, Y100n.

### ATAC-seq mapping and peak calling

We mapped the ATAC-seq reads to the mm10 mouse assembly using bowtie2^34^, with default parameters. We removed duplicate reads with the “MarkDuplicates” function from Picard (http://broadinstitute.github.io/picard/), and subsequently also removed mitochondrial reads using samtools^35^. We then created genotype-specific meta-samples, by merging all the individual bam files corresponding to samples from mice of a given genotype. This yielded one meta-sample for KS1, one for KS2, and one for RT1. For wild-type mice, we created two such meta-samples, one from the wild-type littermates of the KS1 and KS2 cohorts (to which the KS1 and KS2 mutant mice were compared to), and one for the wild-type littermates of the RT1 cohort (to which the RT1 mutant mice were compared to). For each of the 5 resulting meta-samples, we then called peaks using MACS2^36^, with the “keep-dup” parameter equal to “all”.

### ATAC-seq differential analysis

We first defined the set of features to be tested as differential, by unionizing the peaks from all meta-samples. After excluding intervals overlapping ENCODE blacklisted regions^51^, we obtained 78,193 genomic intervals (median size = 690bp, 95^th^ percentile = 1,774, range = 151 - 11363). To verify that these intervals are not likely to be false positives, we compared them to publicly available DNase Hypersensitivity Sites in B cells (CD19+) from the ENCODE project (https://www.encodeproject.org/experiments/ENCSR000CMM/). We converted the DHS coordinates from mm9 to mm10 using liftOver. We then unionized the intervals from the two DHS replicates to create a common set of 112,728 DHS’s. We found that 78,101 of our 78,193 regions (99.88%) overlapped DHS’s, providing strong orthogonal evidence that they represent true B cell regulatory regions.

We then counted the number of reads from each sample that map to each of the 78,193 features, using the featureCounts function from the Rsubread R package^37^, with the following parameters: requireBothEndsMapped = TRUE, countChimericFragments = FALSE, countMultiMappingReads = FALSE, minOverlap = 3. This resulted in a count matrix with rows corresponding to features (the aforementioned genomic intervals), and columns to samples. This count matrix served as input for the differential analysis, which we performed using DESeq2^38^, retaining only features with a median (across samples) count greater than 10. We used Surrogate Variable Analysis^39^ to estimate unobserved confounding variables, and adjusted for those in the differential analysis (without explicitly including other covariates in the model). To derive the list of features overlapping promoters, we first obtained promoter coordinates with the “promoters” function from the EnsDb.Mmusculus.v79 R package, with the parameters “upstream” and “downstream” both equal to 2000. We subsequently restricted to protein-coding transcripts, using the “tx_biotype” filter. The overlapping features were then obtained using the findOverlaps function from the GenomicRanges R package.

### RNA-seq

We spun approximately 100k-500k cells at 300-500g for 5 min at 4’, homogenized in Trizol (Invitrogen 15596018) and stored at −80 until extraction. We extracted and isolated RNA by phase separation using standard protocols followed by purification using the Direct-zol RNA microprep kit (Zymo R2060) with an on-column DNAse step per manufacturer directions. Once purified, we quantified RNA using Quant-iT RiboGreen RNA Assay Kit (ThermoFisher R11490) or Qubit RNA HS Assay Kit (ThermoFisher Q32852), and checked quality by Bioanalyzer with RNA 6000 Pico Kit (Agilent 5067-1513). All samples show high quality RNA with RIN greater than 9. We used 20ng RNA per KS1 & KS2 & matched wild-type sample and 100ng per RT & matched wild-type sample as input to capture mRNA (NEBNext Poly(A) mRNA Magnetic Isolation Module; NEB #E7490) followed by library generation using NEBNext Ultra II Directional RNA Library Prep Kit for Illumina (NEB E7760/E7765) per manufacturer protocols. We determined library size and quality using BioAnalyzer with DNA High Sensitivity reagents (Agilent 5067-4626), and determined concentration using Qubit dsDNA HS Assay Kit (ThermoFisher Q32851) and KAPA Library Quantification Kit for qPCR (KAPA KK4824). We pooled samples and sequenced on Illumina HiSeq4000 using PE flow cells with 100-8-8-100 read length using standard manufacturer protocols. Samples were clustered to aim for 60M reads per sample. Samples were demultiplexed using Illumina pipeline bcl2fastq2 v2.20.

### RNA-seq mapping and differential analysis

We first obtained a fasta file (Mus_musculus.GRCm38.cdna.all.fa.gz) containing all mouse cDNA sequences from Ensembl (http://uswest.ensembl.org/Mus_musculus/Info/Index, version 91, downloaded January 2018). We used this file to build an index and pseudo-map the RNA-seq reads with Salmon (v0.10)^40^. We subsequently imported the resulting transcript quantifications into R to get gene-level counts, using the tximport R package^41^. The differential analysis was then performed with DESeq2, following the same steps as with ATAC-seq.

### Principal Component Analysis

All PCA plots were generated as follows. We first applied a variance stabilizing transformation to the count matrices (either genes-by-samples or genomic-intervals-by-samples), as implemented in the vst function from DESeq2. We then used the resulting matrix to perform the PCA with the plotPCA function.

### Testing for statistically significant overlap between two lists of differential features and identifying the common hits

Our problem is cast in the following setting. Assume we have performed two experiments, each of which involves measuring multiple features (e.g. genes or peaks) in two conditions and performing a differential analysis. The two experiments measure the same set of features. Because the two experiments investigate different biological systems, we don’t expect the set of (true) differential features to be identical. But we are interested in the extent of the overlap between the two sets of features, specifically

a. Is there statistically significant overlap between the two sets of differential features and how big is it?
b. Which features are differential in both lists?

Our approach to these questions is a conditional approach: we ask, does information about the result in experiment 1 affect our interpretation of experiment 2?

We first (arbitrarily) designate one of the two experiments as experiment 1. We test *m* features, and for each feature *i*, we let *X_i_* be a factor with values in {0,1} expressing whether the feature was significantly differential in experiment 1 (*X_i_* = 1), or not (*X_i_* = 0). We are interested in whether the variable *X* = (*X*_1_, …, *X_m_*) is an informative covariate for experiment 2, using terminology from recent work in covariate-powered multiple hypothesis testing^42,43^.

We now consider experiment 2. We split the features into two groups, conditional on the results in experiment 1. Group 1 consists of the features which were found to be differential in experiment 1 (the size of group 1 is *n*) and group 0 consists of the features which were not differential in experiment 1 (of which we have *m* – *n*). We let each group have its own proportion of differential features, that is we introduce parameters *π*_1 | 0_ and *π*_1 | 1_. Let *Y_i_* be the indicator whether the *i*’th feature is differential in experiment 2 or not. Then

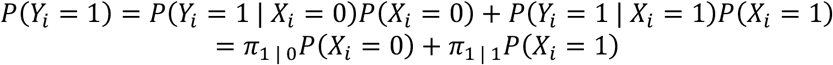

Our null hypothesis is that experiment 1 is not informative about experiment 2, or in other words

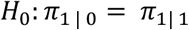

Under this null hypothesis *P*(*Y_i_* = 1) = *π*_1 | 1_. Furthermore, *π*_1 | 1_ should have the same distribution as the proportion of significant features in a random sample of *n* features from experiment 2, which we term 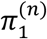.

This gives us the following method for testing *H*_0_:

1. Analyze experiment 1 and decide which features are significantly differential or not.
2. Analyze experiment 2, but only the features which were called differential in experiment 1 to estimate 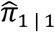.
3. Repeatedly, draw *n* features and estimate 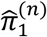 to get a null distribution.

In practice we can estimate 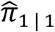 and 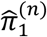 using a number of different methods that produce estimates of the proportion of true null hypotheses (and thus, of the proportion of false null hypotheses, our statistic of interest here) among a set of hypotheses tested. We here used Storey’s method as implemented in the qvalue package in R^44,45^, with the “pi0.method” parameter set to “bootstrap”. This tells us the size of the overlap and the extent to which it is significantly greater than what expected by chance. To estimate which features are in the overlap, we use the “qvalue” function on the features which are significant in the analysis of experiment 1, with the FDR level set to 10%.

### Pairwise comparisons between the disorders

We identified greater overlap between the differentially accessible regions identified in KS1 and KS2, than between the differentially accessible regions identified in either of KS1 or KS2 and RT1. To verify that this is not driven by the fact that KS1 and KS2 were compared against the same wild-type group, we re-estimated the overlap, after first conducting differential analyses where KS1 and KS2 mice were compared to separate wild-type cohorts (8 wild-type mice for the KS1 cohort, and 4 mice for the KS2 cohort, respectively). This again revealed the same picture: 69.5% of differentially accessible regions in KS1 were estimated to be differential in KS2 as well, whereas only 22.8% of differentially accessible regions in RT1 were estimated as differential in KS2.

### Identification of differentially expressed genes with differentially accessible promoter peaks

For **Figure 3b**, we first selected the genes downstream of the top 1000 differentially accessible promoter peaks, the latter being ranked based on their p-values in each disorder. Out of these genes, we retained those differentially expressed using the “qvalue” function from the qvalue R package, with the gene p-values as input and the “fdr.level” parameter set to 0.1. In cases where there were more than one peaks in the same promoter, we calculated the median logFC across these peaks. For **Figure 3a**, we slid the threshold for the top differentially accessible promoter peaks in each disorder from 1000 to 5000, and estimated the proportion of differentially expressed downstream genes used the “pi0est” function from the qvalue R package with the “pi0.method” parameter set to “bootstrap”.

Finally, for **Figure 4h** we employed the analogous procedure to **Figure 3a** in order to estimate the proportion of differentially accessible peaks in promoters of differentially expressed genes for different thresholds.

### Reactome pathway analysis

We used the goseq R package^46^ to perform pathway analyses for the commonly disrupted genes, based on Reactome pathways^47^. As our assayed gene set, we used the set of all genes included in all three differential expression analyses, or the set of all genes that had at least one promoter peak included in all three differential accessibility analysis. As our differential gene set, we used the set of genes commonly differentially expressed across the three MDEMs, or the set of genes with at least one commonly differentially accessible promoter peak. The top 20 enriched pathways are provided in **Supplemental Tables 9** and **10**, respectively.

### Transcription Factor motifs

We obtained a bed file (mm10.archetype_motifs.v1.0.bed) containing the genomic positions of 233 non-redundant TF motifs, from https://www.vierstra.org/resources/motif_clustering^48^. We then restricted to motifs that had at least 1 base overlapping our set of unionized B cell peaks (see sections **ATAC-seq mapping and peak calling** and **ATAC-seq differential analysis**). Subsequently, we tested each motif for enrichment as described in the results section using the fisher.test function in R.

### Gene catalogs

#### Transcription Factors

We obtained a list of 1,254 genes encoding for human TFs from Barrera et al., 2016.^49^ We the used the biomaRt R package to obtain the mouse orthologs of these TF genes, with the ENSEMBL ids as our filter. We only retained high-confidence orthologs (“mmusculus_homolog_orthology_confidence” equal to 1). Finally, we restricted to TFs included in all three differential analyses (KS1 vs WT, KS2 vs WT, and RT1 vs WT).

#### IgA deficiency genes

We used the Mammalian Phenotype Browser on the Mouse Genome Informatics database^50^ to obtain a catalog of genes known to lead to IgA deficiency when individually knocked-out. Specifically, we used “decreased IgA level” as the phenotype term and then obtained all the resulting genes, regardless of the genetic background. In cases of double knockouts, we included both genes.

### ELISA for serum IgA levels

We performed ELISAs on serum IgA from peripheral blood samples as previously described^13^.

## Code availability

All code for the analyses in this manuscript is available at https://github.com/hansenlab/mdem_overlap

## Data availability

The ATAC- and RNA-seq data generated for this study are available in GEO (accession GSE162181).

## Acknowledgements

H.T.B. and T.R.L. are supported by a grant from the Louma G. Foundation. H.T.B. is also supported by grants from the Icelandic Research Fund (#195835-051, #206806-051) and the Icelandic Technology Development Fund (#2010588-0611). K.D.H, H.T.B and LB were partly supported by the 2016 Discovery Award from Johns Hopkins University. L.B. was partly supported by the Maryland Genetics, Epidemiology and Medicine (MD-GEM) training program, funded by the Burroughs-Wellcome Fund. Research reported in this publication was supported by the National Institute of General Medical Sciences of the National Institutes of Health under award number R01GM121459. **Figure 1d** was in part created using Biorender; this software has a restrictive license which includes a requirement to be mentioned in the acknowledgements.

## Supplemental Figures

**Supplemental Figure 1.**
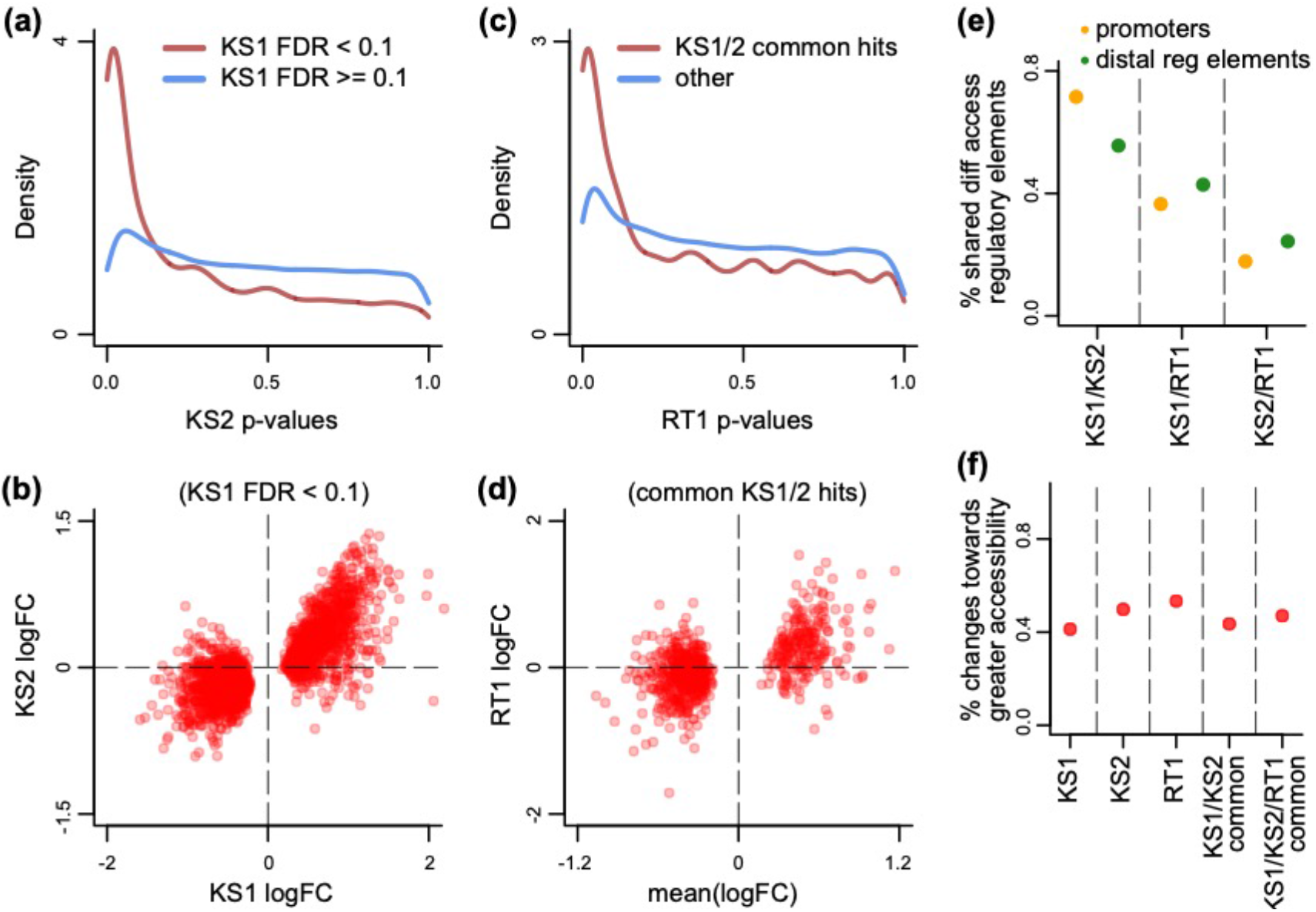
Evaluating the overlap between the differentially accessible distal regulatory elements in B cells in Kabuki type 1, Kabuki type 2, and Rubinstein-Taybi syndromes. **a**) The distribution of p-values from the KS2 vs wild-type differential accessibility analysis for peaks at distal regulatory elements (defined as peaks not within +/- 2kb from the TSS), stratified according to whether the same elements are significantly differentially accessible in KS1 (FDR < 0.1; red curve), or not (FDR >= 0.1; blue curve). **b**) Scatterplot of log2(fold changes) from the KS1 vs wild-type (x-axis) distal regulatory element differential accessibility analysis against the corresponding log2(fold changes) from the KS2 vs wild-type analysis (y-axis). Each point corresponds to a peak. Shown are only peaks that are differentially accessible in KS1 (FDR < 0.1). **c**) The distribution of p-values from the RT1 vs wild-type differential accessibility analysis for distal regulatory element peaks, stratified according to whether the same peaks are commonly differentially accessible in KS1 and KS2 (FDR < 0.1, see Methods; red curve), or not (blue curve). **d**) Scatterplot of log2(fold changes) from the RT1 vs wild-type (x-axis) differential accessibility analysis, against the mean log2(fold change) from the KS1 vs wild-type and KS2 vs wild-type analyses. Each point corresponds to a peak. Shown are only peaks that are commonly differentially accessible in KS1 and KS2 (FDR < 0.1). **e**) The pairwise overlap between the differentially accessible peaks (promoters or distal regulatory elements) in the three MDEMs. **f**) The proportion of differentially accessible distal regulatory elements that show increased accessibility in the mutant vs the wild-type mice.

## Supplemental Tables

We provide the following Supplemental Tables:

1. Coordinates of promoter peaks commonly differentially accessible in KS1 and KS2, along with the corresponding logFC changes.
2. Coordinates of promoter peaks commonly differentially accessible in KS1, KS2, and RT1, along with the corresponding logFC changes.
3. Coordinates of distal regulatory element peaks commonly differentially accessible in KS1 and KS2, along with the corresponding logFC changes.
4. Coordinates of distal regulatory element peaks commonly differentially accessible in KS1, KS2, and RT1, along with the corresponding logFC changes.
5. Differentially expressed genes downstream of differentially accessible promoter peaks, along with the corresponding p-values and logFC changes.
6. Genes commonly differentially expressed in KS1 and KS2, along with the corresponding logFC changes.
7. Genes commonly differentially expressed in KS1, KS2, and RT1, along with the corresponding logFC changes.
8. Transcription factor motifs enriched in peaks found in promoters of differentially expressed genes.
9. Top 20 Reactome enriched pathways, using genes commonly differentially expressed in KS1, KS2, and RT1.
10. Top 20 Reactome enriched pathways, using genes with commonly differentially accessible promoters in KS1, KS2, and RT1.

